# Interactomics-driven discovery of alkaloid biosynthetic pathways in kratom

**DOI:** 10.64898/2026.05.01.722234

**Authors:** Bo Yang, Samuel Kemiji, Jianing Han, Dong Oh Han, Yinan Wu, Chang Liu, Frank C. Schroeder, Sijin Li

## Abstract

Plant monoterpene indole alkaloids (MIAs) exhibit important pharmacological activities, yet understanding of their biosyntheses remains incomplete. Since protein-protein interactions (PPIs) represent a conserved regulatory mechanism in MIA-producing plants, we developed a large-scale, yeast-based screening pipeline to profile PPIs of a key enzyme, strictosidine β-D-glucosidase (SGD) from *Mitragyna speciosa* (kratom). This screen identified six novel medium-chain dehydrogenases/reductases (MDRs) as high-confidence interaction partners of SGD. Biochemical characterization revealed that all six MsMDRs produce an MIA we named charlamine by acting directly on the reactive strictosidine aglycone intermediate, preventing its spontaneous rearrangement and establishing a functional rationale for SGD–MDR interaction. One MsMDR additionally catalyzed the reduction of vallesiachotamine, derived from the spontaneous rearrangement of strictosidine aglycone, to another previously unreported MIA, vallesiachotaminol. Parallel transcriptomics and genomics analyses uncovered a biosynthetic gene cluster containing a dihydrocorynantheine aldehyde esterase, functioning downstream of MsMDRs. Collectively, these findings demonstrate the utility of interactomics-driven plant pathway discovery.

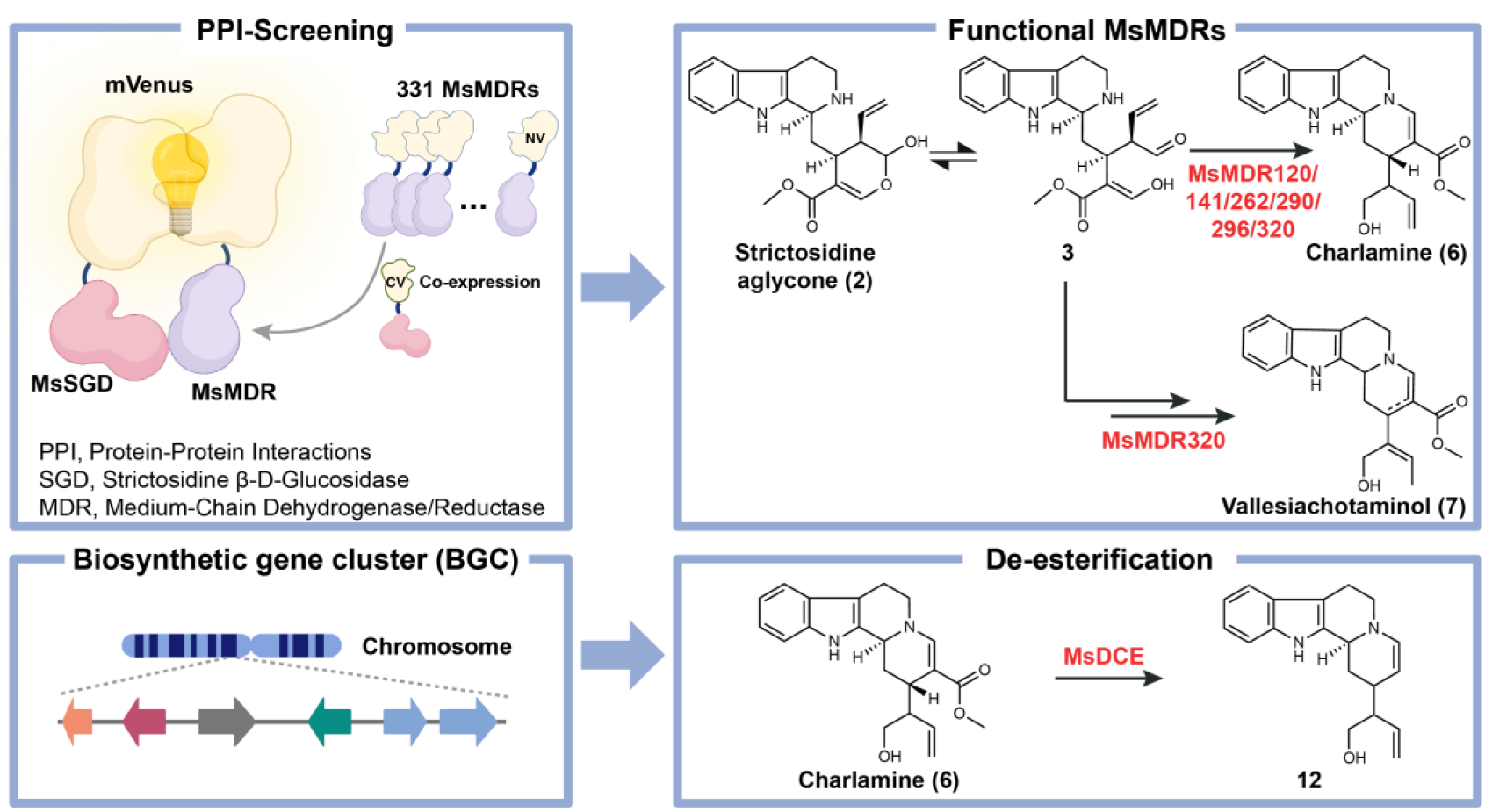

## Main

Monoterpene indole alkaloids (MIAs) are an important family of plant natural products that exhibit complex biochemistry controlled by sophisticated regulatory mechanisms^1–3^. MIA biosynthesis in planta relies on transient protein-protein interactions (PPIs)^4^ that assemble individual enzymes into dynamic enzyme complexes to organize and regulate diverse MIA pathway branches at the post-translational level^5^. Binary PPIs between the gateway enzyme, strictosidine beta-D-glucosidase (SGD), in MIA biosynthesis, and its downstream enzymes, medium-chain dehydrogenases/reductases (MDRs), have been reported in multiple MIA-producers, such as *Catharanthus roseus* (Madagascar periwinkle)^6–8^ that produces the anticancer agent vinblastine and *Mitragyna speciosa* (kratom)^9^ that produces analgesic mitragynine^10^. These SGD-MDR PPIs have been proposed to serve as a conserved mechanism that prevents the diffusion of the toxic strictosidine aglycone (**2a**) produced by SGD from strictosidine (**1**) by rapidly converting it to more stable downstream MIAs via MDRs. Our prior study leveraged the PPIs between putative MDRs and SGD as a unique pattern for pathway prediction and combined it with transcriptomics analysis to discover novel MDRs from kratom. Combined yeast-(*Saccharomyces cerevisiae*) based PPI identification and biochemical characterization led to the identification of four functional MsMDRs that interact with MsSGD, including tetrahydroalstonine synthase (MsTHAS), heteroyohimbine synthase (MsHYS), dihydrocorynantheine synthase (MsDCS), and a novel MDR with a hitherto uncharacterized product (MsMDR4)^9^. These MDRs, along with geissoschizine synthase (GS) derived from *C. roseus*^8^, represent the only five types of MDRs known to interact with SGD in MIA-producing plants.

In this work, we globally profiled PPIs between MsSGD and all kratom proteins putatively annotated as MsMDRs-like enzymes to gain further structural insights into kratom MIA biosynthesis, including the roles of novel MsMDRs. We developed a large-scale PPI screening pipeline in yeast using the bimolecular fluorescence complementation (BiFC)^11^ method and flow cytometry to enable high-throughput screening. Using this yeast-based platform, we screened 331 kratom genes putatively annotated as MsMDRs or related enzymes^12^. The top 60 interactor candidates were then screened biochemically in yeast, yielding nine functional MsMDRs, including three previously characterized interacting MsMDRs, validating our screening platform, as well as six MsMDRs that all catalyze formation of a previously uncharacterized MIA, charlamine (**6**), as a major product. Among the six MsMDRs we selected MsMDR290 for further biochemical characterization and demonstrated that the biosynthesis of **6** involved the reduction of a transient aldehyde (**3**) to the corresponding alcohol (**4**), followed by spontaneous cyclization and dehydration (**5**), resulting in the formation of the more stable 1,2-dihydropyridine in **6**. In addition, we found that one of the six MsMDRs, MsMDR320, can reduce reactive α, β-unsaturated aldehydes (**8a/b**) derived from hydrolysis of strictosidine to the corresponding alcohols, which we named vallesiachotaminol (**7a** and **7b**). Metabolomic analysis of kratom samples demonstrated that **6** and **7** are also produced in the plant. Leveraging the novel MsMDRs as baits in transcriptomic and genomic analyses, we identified an MsMDR-rich biosynthetic gene cluster (BGC) encoding putative novel MsMDRs, previously characterized MsDCSs, and a homolog of dihydrocorynantheine aldehyde esterase (DCE) identified from *Cinchona pubescens*^13^. We show that this enzyme, named MsDCE, catalyzes the de-esterification of both 20*R*/*S*-dihydrocorynantheines (**10**) produced by MsDCS and **6** produced by the newly discovered MsMDRs, revealing two MIA pathway branches and associated products (**11** and **12**) (Fig. 1). This work demonstrates the potential of interactomics analysis for plant natural product (PNP) pathway discovery, which can capture the spatial organization of pathway enzymes, complementing conventional prediction methods based on transcriptomics and genomics to accelerate the PNP pathway discovery.

**Fig. 1:**
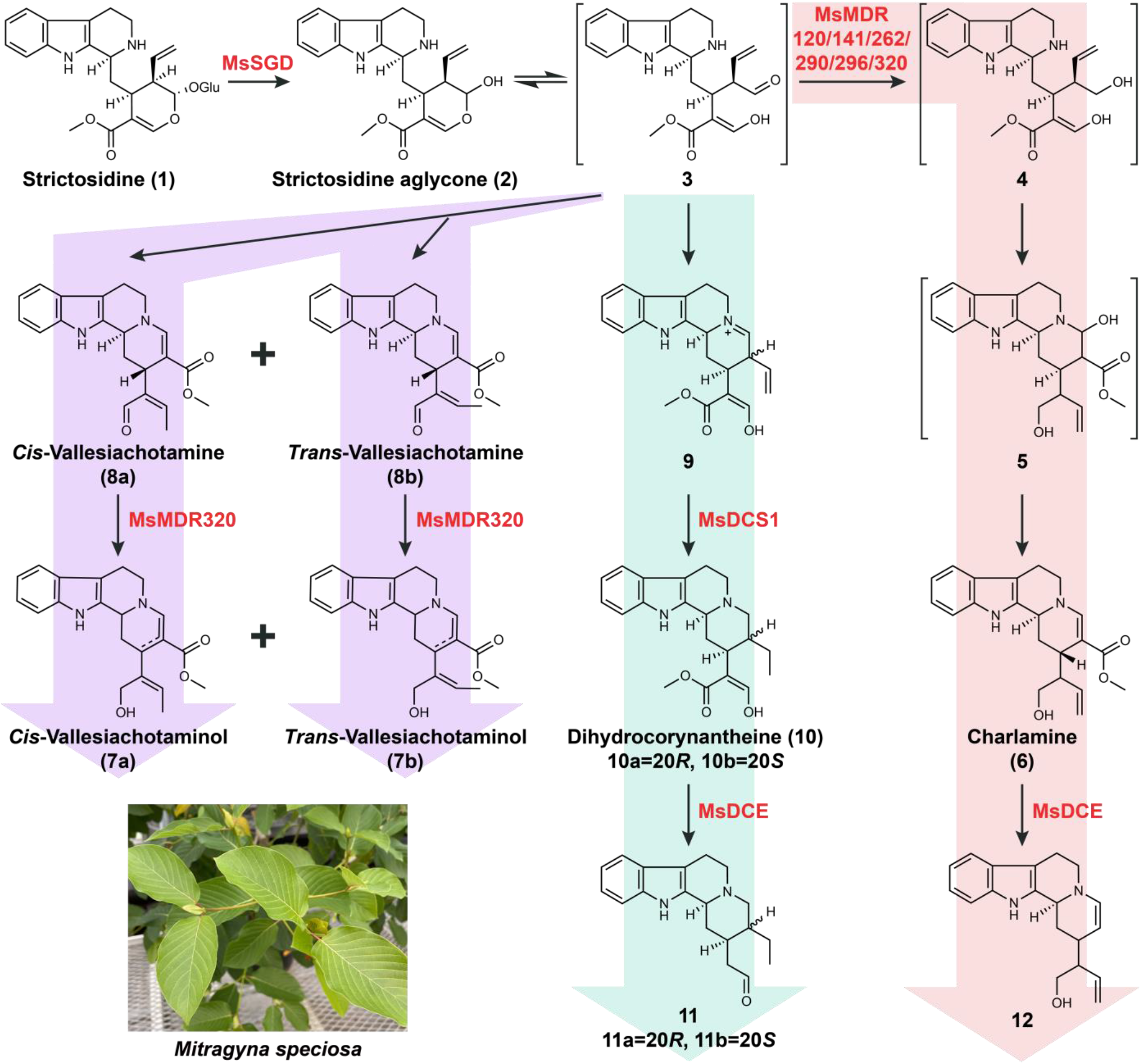
Scheme of MIA biosynthetic pathways discovered in this work. Novel MsMDRs, represented by MsMDR290 or MsMDR320, act directly on the transient aldehyde (**3**) form of strictosidine aglycone, producing alcohol (**4**), followed by spontaneous cyclization and dehydration (**5**), resulting in the formation of charlamine (**6**). MsMDR320 also catalyzes reduction of the aldehyde group of vallesiachotamine (**8**) to the corresponding alcohol, vallesiachotaminol (**7**). In contrast, MsDCS1 acts on the rearranged form of strictosidine aglycone (**9**) to yield dihydrocorynantheine (**10**). Newly discovered MsDCE accepts both **6** and dihydrocorynantheine (**10**) produced by characterized MsDCS1 to produce de-esterification products **12** and **11**, respectively.

## Results

### A yeast-based PPI screening platform identified 60 MsMDR candidates

To profile putative MsSGD-MsMDR PPIs in yeast, we employed the BiFC method, which provides high fidelity for capturing transient PPIs by the complementation of two mVenus split fragments fused to MsSGD and the putative MsMDRs, enabling the use of flow cytometry as a high-throughput method to identify interacting MsMDRs. Similar to our reported design^9^, we fused the C-terminal mVenus split fragment (CV) to the N-terminus of MsSGD (CV-MsSGD) as bait. Conversely, the N-terminal mVenus split fragment (NV) was fused to the C-terminus of putative MsMDRs as prey. This design maintained the C-terminal nuclear localization sequence (NLS)^9^ of MsSGD and minimized the impact on MsMDR localization in yeast, as demonstrated in our previous study. We selected multi-copy plasmids rather than chromosomal integration in yeast to express the candidate genes effectively, thereby avoiding the potential loss of weak interactors.

First, we developed a host yeast strain containing a multi-copy plasmid expressing CV-MsSGD as bait. Next we identified a total of 331 putative MsMDRs from the published kratom transcriptomic data^14^ to be used as prey, based on annotations as MDRs, cinnamyl alcohol dehydrogenases (CADs), or ADHs. This list included the previously characterized MsHYS, MsTHAS, and MsDCS1. Each MsMDR gene was cloned from kratom cDNA in a plate-based high-throughput manner and co-transformed with a pre-established linearized vector containing the NV fragment into the host strain, yielding 331 yeasts each co-expressing one pair of CV-MsSGD and MsMDR-NV. After transformation, yeast constructs were spread on synthetic dropout (SD) plates. Colonies were then cultured for two days and analyzed by flow cytometry (Fig. 2a).

**Fig. 2:**
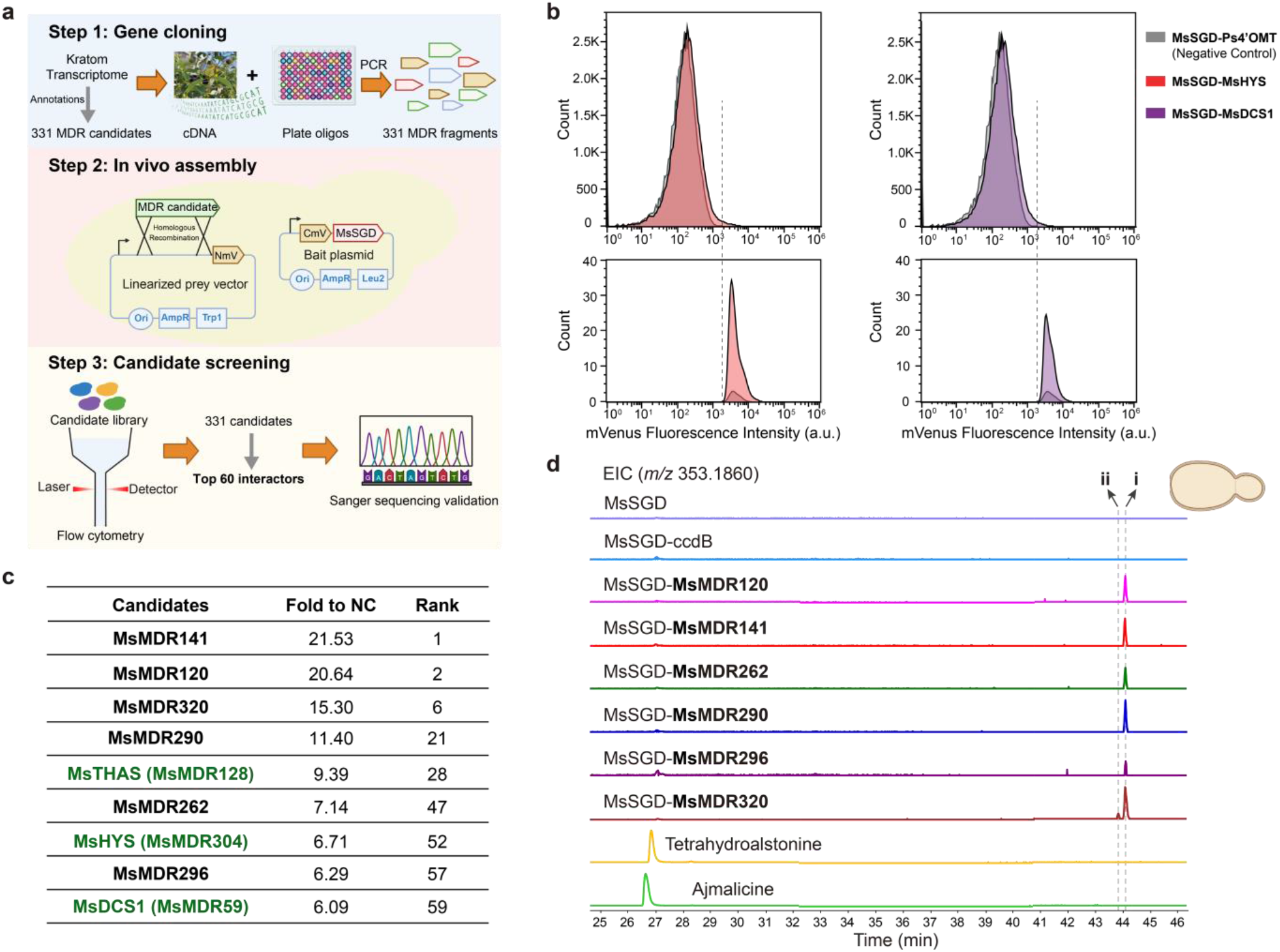
Yeast-based large-scale PPI screening and biochemical characterization identified novel MsMDRs. a) Scheme of a yeast-based BiFC pipeline for large-scale PPI screening using flow cytometry.b)Positive interactors, including MsHYS and MsDCS1, exhibited 6.9-fold and 4.9-fold of cells compared to the negative control with a cutoff at 5,000 a.u. in a preliminary test, setting the threshold for screening.c)The rank of fluorescence intensity of nine MsMDRs among all candidates. d) LC-Q-TOF analysis of yeasts co-expressing MsMDR and MsSGD identified six novel functional MsMDRs. EIC of *m/z* [M+H]^+^=353.1860 (20 ppm) led to two unique peaks, **i** (identified in all six MsMDR-expressing yeasts) and **ii** (only identified in the presence of MsMDR320).

To identify true interactors, we included characterized MsHYS and MsDCS1 as positive controls and the non-interacting Ps4’OMT, an unrelated methyltransferase from *Papaver somniferum*^15^, as a negative control to establish a screening threshold. Initial tests using flow cytometry and plate reader assays showed that transient PPIs between MsSGD and MsMDR did not result in a significant difference in average fluorescence, which thus cannot be used as a criterion for selecting positive interactors. However, the known interactors consistently generated a unique, high-fluorescence sub-population in flow cytometry analysis (Fig. 2b), in alignment with prior microscopic analysis^9^. Fig. 2b demonstrates the significantly increased high-fluorescence sub-population in case of MsHYS and MsDCS1. Based on a fluorescence intensity cutoff at 5000 a.u., only 0.074% cells in the negative control exhibited high fluorescence, while 0.510% cells expressing MsHYS and 0.359% cells expressing MsDCS1 fell within the high-fluorescence group, corresponding to 6.9- and 4.9-fold increases over control, respectively. The fold increase of this high-fluorescence sub-population over the negative control was then chosen as the criterion for identifying positive interactors. Across multiple repeats, the fold increase for the two pre-established positive controls was consistently above six-fold. Therefore, all MsMDRs with greater than six-fold increase relative to the negative control were considered PPI-positive, leading to identification of a total of 60 MsMDRs as positive interactors (Supplementary Table 1).

### Biochemical screening and PPI assays in yeast identified six functional MsSGD-interacting MsMDRs

We screened for the production of new MIAs by expressing the 60 identified candidate MsMDRs in yeast, followed by analysis via HPLC coupled to high-resolution mass spectrometry (HRMS). MsSGD and each MsMDR were cloned into separate multi-copy plasmids and transformed into *S. cerevisiae* CEN.PK2-1D. Two negative controls were included: one expressing only MsSGD and another co-expressing MsSGD and the unrelated gene ccdB, replacing the MsMDR cassette. Approximately 0.15 mM strictosidine, generated by *in vitro* enzymatic synthesis, was added at the start of fermentation, and its consumption and appearance of putative products after 72 h of fermentation were probed by HPLC-HRMS. Nine of the 60 MsMDR candidates, including the three previously characterized interactors, MsHYS, MsTHAS, and MsDCS1, were found to be active (Fig. 2c). The enzymatic activities of MsHYS, MsTHAS, and MsDCS1 were consistent with previous observations^9^ (Supplementary Fig. 1). More importantly, the six remaining MsMDRs (Fig. 2c), all produced one major common MS feature with a mass/charge ratio (*m/z*) of 353.1860 (peak **i**, Fig. 2d) and a retention time that did not match those of standards of known isobaric MIAs (i.e., ajmalicine and tetrahydroalstonine), indicating a distinct structure. In addition to peak **i**, MsMDR320 produced a slightly earlier eluting MS feature at *m/z* of 353.1860 (peak **ii**, Fig. 2d), while the other MsMDR candidates did not yield additional unique MS features based on untargeted metabolomics analysis.

BiFC and protein localization assays further confirmed positive PPI between MsSGD and the six newly identified functional MsMDRs. All BiFC fluorescence signals localized to the yeast nucleus (Extended Data Fig. 1). In the protein localization assay, the six MsMDRs localized to the cytoplasm when expressed alone in yeast (Extended Data Fig. 2a) but relocalized to the nucleus upon co-expression with MsSGD (Extended Data Fig. 2b).

Phylogenetic analysis of the 60 candidates, together with previously characterized MsMDRs, showed that five of the novel MsMDRs clustered, whereas MsMDR296 formed a relatively more distant branch (Supplementary Fig. 2). Subsequent structure alignment (Supplementary Fig. 3) and biochemical analysis (Supplementary Fig. 4) revealed that MsMDR296 also shared high structural and functional similarities with dihydroflavonol 4-reductase. Combined with its lowest activity in producing **i** (Fig. 2d), MsMDR296 might be a bifunctional enzyme acting on both flavonols and MIAs. For subsequent product characterization and functional analysis, we focused on MsMDR290, which showed the highest activity among the six MsMDR candidates tested in yeast, and MsMDR320, which produced an additional previously uncharacterized MS feature.

**Fig. 3:**
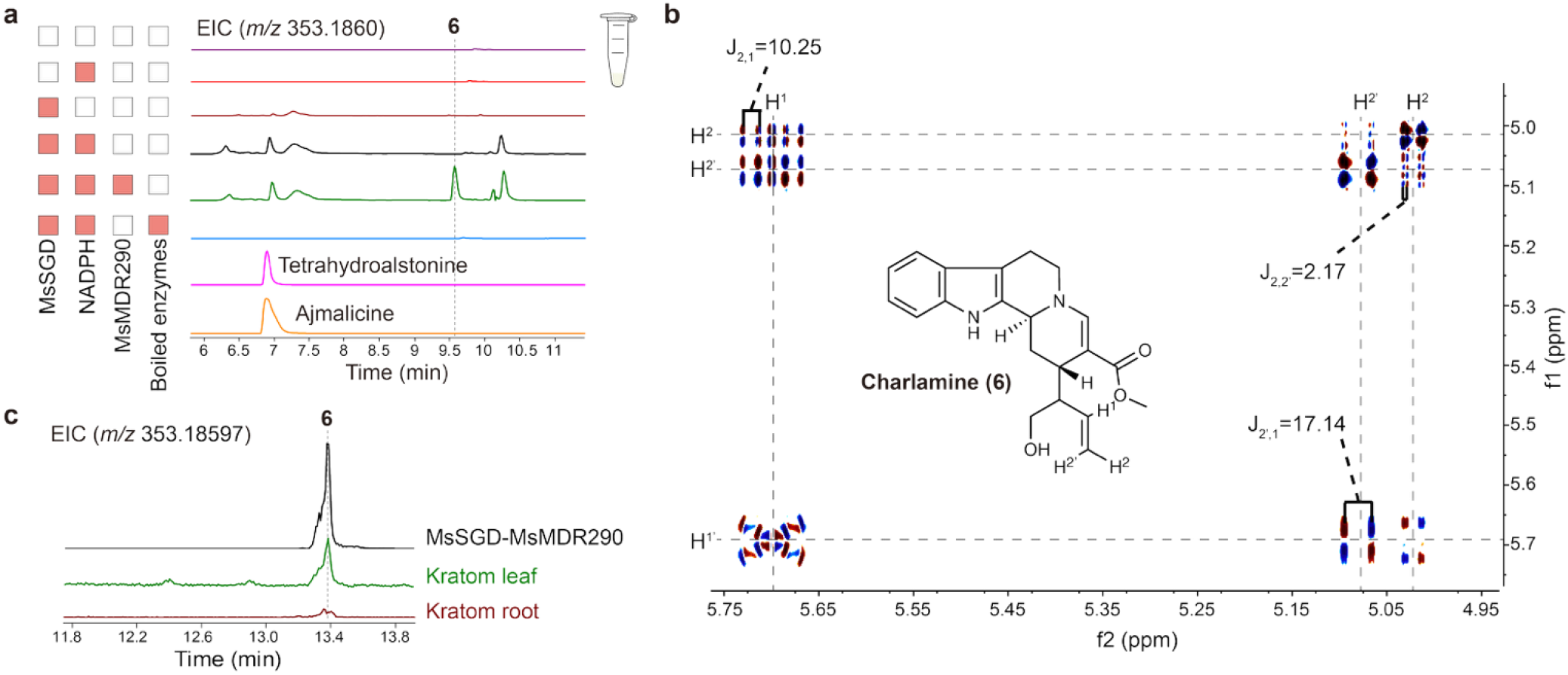
Identification of charlamine (6) as major *in-vitro* product of MsMDR290 and its detection via kratom metabolomics analysis. a) LC-Q-TOF analysis (20 ppm) identified **6** synthesized from strictosidine, catalyzed by MsSGD and MsMDR290 in in vitro assays. Negative controls were performed by replacing soluble MsSGD and MsMDR290 with boiled proteins. b) Structure elucidation of charlamine (**6)** in methanol-*d*_*4*_using dqfCOSY NMR. Spectrum showing correlations with the terminal alkene (2.3-5.8 ppm, 600 MHz). c) Metabolite profiling of kratom plants using Orbitrap (10 ppm) revealed the accumulation of **6** in leaves and roots.

**Fig. 4:**
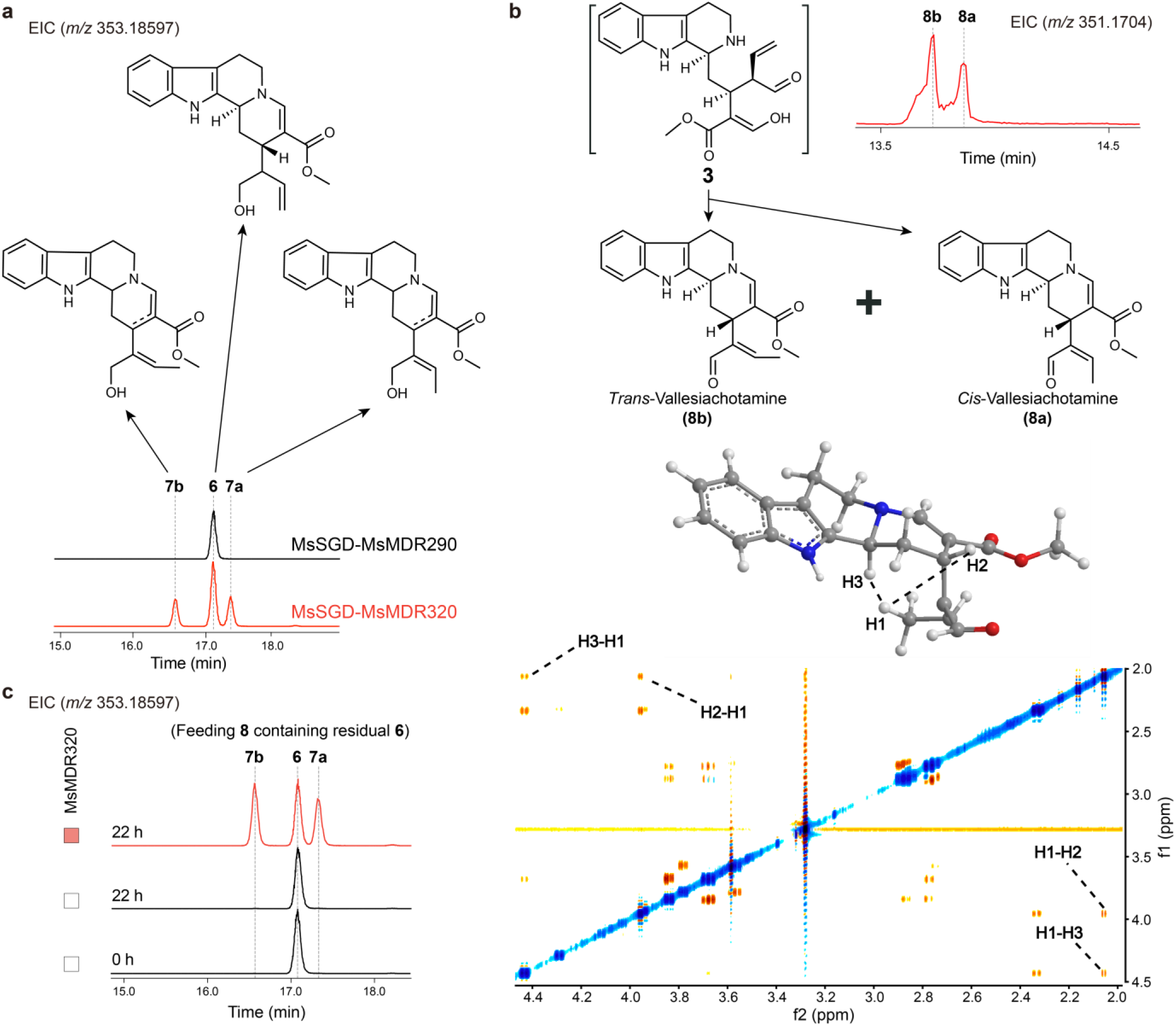
Discovery and characterization of vallesiachotaminol (7), a reduction product of vallesiachotamine by MsMDR320. a) LC-Q-Orbitrap analysis (10 ppm) of in vitro assay samples. Vallesiachotaminol (**7**) was identified by LC-MS following in vitro conversion of strictosidine catalyzed by MsSGD and MsMDR320 (Red). b) An overnight hydrolysis product of strictosidine with *m/z*=351.1704 was detected and identified as vallesiachotamine (**8**) by NMR analysis. NOESY spectrum (600 MHz) of vallesiachotamine (**8)** produced in vitro in methanol-*d*_*4*_ indicated a *cis*:*trans* ratio of 1:3. c) In vitro assays showing that MsMDR320 could convert vallesiachotamine (**8**) to the corresponding alcohol, **7**. The isolated **8** used as the substrate contained residual **6** from the purification process.

### MsMDR290 acts on the aldehyde form of strictosidine aglycone

MsMDR290 and MsSGD were expressed in *E. coli* and purified for *in vitro* assays using strictosidine as the substrate. Strictosidine, MsMDR290, and MsSGD were incubated at 30°C for 22 hours. Incubation with MsMDR290 produced one major peak at *m/z* 353.1860 (peak **i**) in addition to several other isobaric features that were also observed when incubating with MsSGD and NADPH only (Fig. 3a). Peak **i** was isolated via silica gel and normal-phase column chromatography and then characterized using a suite of 2D NMR spectra (Fig. 3b, Supplementary Table 2, and Supplementary Fig. 5). NMR spectroscopic analysis revealed a typical MIA ring system identical to that of the known vallesiachotamine, distinguished by a hydroxybutenyl substituent featuring a terminal double bond (Fig. 3b), as in the parent compound strictosidine. This is in contrast to the internal double bond in vallesiachotamine, which likely results from non-enzymatic double bond migration via tautomerization driven by the greater stability of conjugated systems. The product of MsMDR290, named charlamine (**6**), was also detected in yeast co-expressing MsSGD and MsMDR290 (Supplementary Figs. 6 and 7) as well as in kratom leaves and roots, particularly enriched in leaves (Fig. 3c), as confirmed by comparison of retention times and MS/MS spectra (Supplementary Fig. 8).

**Fig. 5:**
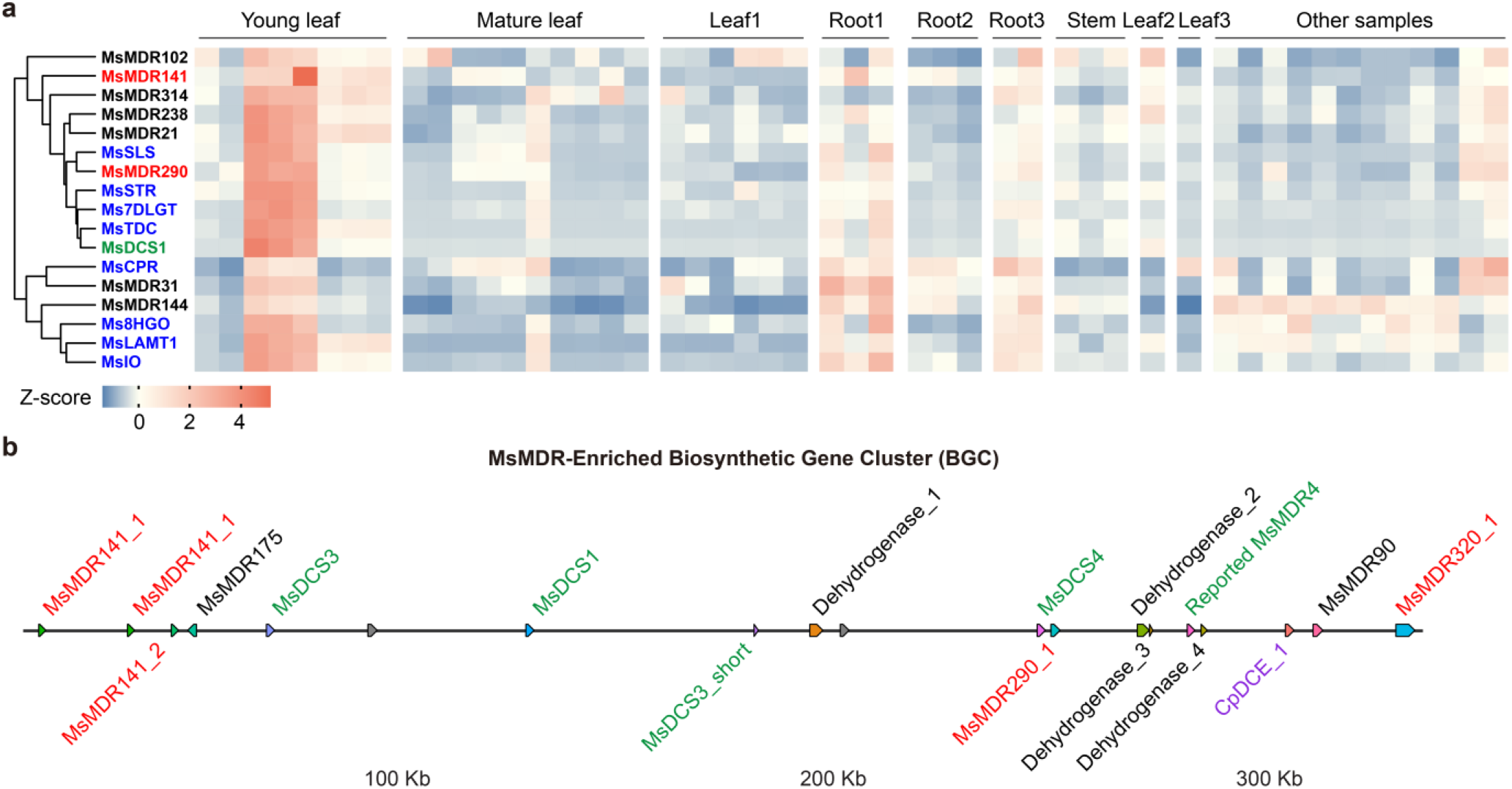
Pathway discovery by transcriptomics and genomic analysis using MsMDRs as baits. a) Transcriptional profiles of 60 positive interactors and the upstream MIA-pathway genes across 49 published kratom transcriptomes were analyzed, and genes exhibiting co-expression patterns are shown. Each column represents a biological replicate, and each row corresponds to a distinct gene. Expression levels are shown as Z-scores calculated from log_2_-normalized TPM values, ranging from −1.4 (blue, lower expression) to +5.1 (red, higher expression). Among the 60 putative interacting MsMDRs, three functional and interacting MsMDRs, such as MsMDR141, MsMDR290, and MsDCS1, were highly expressed in roots from samples with high mitragynine levels (Root 1) and young leaves. Other genes highly expressed include MsSLS, MsSTR, Ms7DLGT, MsTDC, MsCPR, Ms8HGO, MsLAMT1, and MsIO in the strictosidine biosynthetic pathway. Sample descriptors include: Root 1 (high-mitragynine), Root 2 (low-mitragynine), Leaf 2 (leaf bract), and Leaf 3 (leaf after wounding). b) Genomic mapping of the three co-expressed MsMDRs identified a 320-kb biosynthetic gene cluster in Tig00011620 scaffold. Green: previously reported MsMDRs, including MsDCS1, 3, 4, a truncated version of MsDCS3, and MsMDR4; Red: newly discovered functional MsMDRs, including MsMDR141 (three isogenes), MsMDR290, and MsMDR320. Black: putative interacting MsMDRs that did not exhibit activity in biochemical screening. Abbreviations: MDR, medium-chain dehydrogenase/reductase; SLS, secologanin synthase; STR, strictosidine synthase; 7-DLGT, 7-deoxyloganetic acid-O-glucosyl transferase; TDC, tryptophan decarboxylase; DCS, dihydrocorynantheine synthase; CPR, NADPH-cytochrome P450 reductase; 8-HGO, 8-hydroxygeraniol oxidoreductase; LAMT, loganic acid O-methyltransferase; IO, 7-deoxyloganetic acid synthase/iridoid oxidase.

**Fig. 6:**
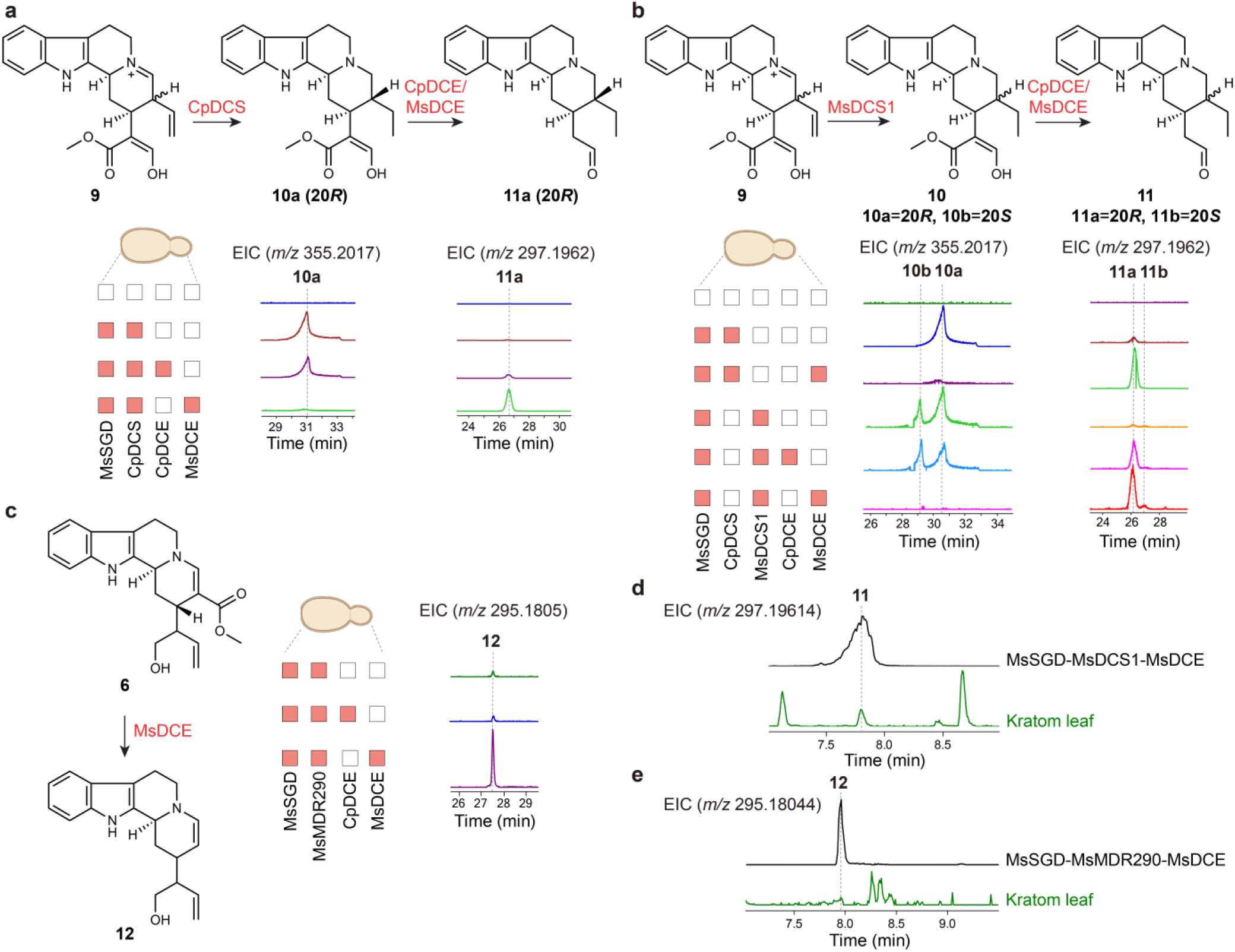
Functional characterization of MsDCE and MsMDRs leading to new downstream pathway branches. a) Reconstitution of the CpDCS-DCE pathway in yeast. Co-expression of MsSGD, CpDCS, and MsDCE converted **10a** to **11a** with substantially higher efficiency than CpDCE. Measured by Q-TOF (20 ppm). b) Reconstitution of the MsDCS1-DCE pathway in yeast. Co-expression of MsSGD, MsDCS1, and MsDCE converted both **10a** and **10b** to **11a** and **11b**. Measured by Q-TOF (20 ppm). c) Yeast co-expressing MsSGD, MsMDR290, and MsDCE produced **12**. Measured by Q-TOF (20 ppm). d) Detection of **11** in kratom leaf, showing alignment with the yeast-expressed MsSGD-MsDCS1-MsDCE products. Measured by Orbitrap (10 ppm). e) Identification of **12** in kratom leaves, matching the MsSGD-MsMDR290-MsDCE reaction product detected in yeast. Measured by Orbitrap (10 ppm).

PPIs between MDRs and SGD have long been postulated to be crucial for avoiding the leakage of toxic intermediates produced by SGD (e.g., **2** and **3**) and directing metabolic flux toward diverse downstream MIAs. The substrates of previously reported interacting MsMDRs, such as MsTHAS, MsHYS, and MsDCS1, are the more stable aglycone isomers (e.g., **8** or **9** in Fig. 1) that derive from the nucleophilic attack of the aliphatic nitrogen onto either the free aldehyde or the enol-masked aldehyde in cytotoxic **3**. Depending on the structure of the resulting products, these known MsMDRs catalyze either 1,2- or 1,4-iminium reduction to form heteroyohimbine- or corynantheine-type MIAs. In contrast, the newly-discovered MsMDRs may act before this iminium or enamine formation occurs, reducing the reactive enol-aldehyde **3** to enol-alcohol **4**, leaving only the more stable enol-masked aldehyde for the nucleophilic nitrogen to react with, which then yields charlamine (**6**), that we here identified (Fig. 1). To enable the swift reduction of the non-conjugated aldehyde in **3**, MsMDRs need to bind the highly reactive **3** before nucleophilic attack of the nitrogen occurs, or the double bond can isomerize to form a conjugated aldehyde as in vallesiachotamine, further supporting a functional rationale for the observed PPIs.

### MsMDR320 catalyzes aldehyde reduction in vallesiachotamine

In addition to peak **i**, corresponding to charlamine (**6**), yeast-based and in vitro assays with MsMDR320 produced a later-eluting isobaric peak **ii** (Fig. 2d and Supplementary Fig. 9), whose MS/MS spectrum showed an intense fragment resulting from water loss, which is largely absent from the MS/MS spectrum of **6**. Notably, the MS/MS spectrum of peak **i** in MsMDR320 samples also showed some water loss, suggesting it may consist of **6** and yet another isobaric isomer that fragments in a manner similar to that of peak **ii**. In fact, variation of chromatographic conditions allowed to resolve peak **i** in MsMDR320 samples into two peaks, representing **6** and a second species whose MS/MS spectrum closely resembled that of peak **ii**, suggesting that, in addition to **6**, MsMDR320 produces two closely related isomers. Detailed analysis of their MS/MS spectra suggested that these two additional isomers may represent the reduced products of the known vallesiachotamine (**8**), which we named vallesiachotaminols **7a** and **7b** (Fig. 1, Fig. 4a, and Supplementary Fig. 10)^16^. Consistent with the idea that MsMDR320 acts on vallesiachotamine (**8**), hydrolysis of strictosidine with SGD in vitro produced two MS features at *m/z* 351.1704, which were identified as a 1:3 mixture of *cis*- and *trans*-vallesiachotamine (**8a** and **8b**)^16^ based on 2D NMR spectroscopic analysis (Fig. 4b, Supplementary Tables 3 and 4, and Supplementary Fig. 11). Previous studies identified **8a/b** from *Alstonia scholaris*^17^ and *C. roseus*^18^, and **8b** from *Anthocephalus chinensis*^19^, although the biosynthetic pathway was not elucidated. Our in vitro assay revealed that **8** can be formed via irreversible isomerization of **3** derived from strictosidine hydrolysis.

To further characterize MsMDR320’s function in converting vallesiacthotamine (**8**) to vallesiachotaminol (**7**), we used a sample of **8** isolated from MsMDR290 reaction mixtures as the substrate for in vitro assays with MsMDR320. Note that **8** used for this assay contained a small amount of **6**. Analysis of the reaction mixture containing the isolated **8** and MsMDR320, **7a** and **7b** were generated after 22 h of incubation, whereas the control reaction without MsMDR320 did not produce any **7**. Meanwhile, the concentrations of the impurity **6** remained unchanged in both control and the assay with MsMDR320 (Fig. 4c). The position of the cyclic double bond in **7** could not be determined from MS/MS analysis and is therefore depicted as a dotted bond (Fig. 4). **8a, 8b**, and trace amounts of **7** were also identified from kratom stems and leaves (Supplementary Figs. 12 and 13). Taken together, we concluded that **7** and **6** belonged to two competing branches of MIA metabolism in kratom, and that MsMDR320 can act on both the aldehyde form of strictosidine aglycone (**3**) and vallesiachotamine (**8**).

### Transcriptomics and genomics identified a BGC containing MsMDRs and MsDCE

To identify potential enzymes downstream of the newly characterized MsMDRs, we performed hierarchical clustering to examine co-expression patterns among upstream MIA-pathway genes and the 60 positive interactors across 49 published kratom transcriptomes^9,14,20–22^. Genes that are highly co-expressed in young leaves are shown in Fig. 5a. Notably, three functional MsSGD-interactors, including MsMDR290, MsMDR141, and MsDCS1, clustered with multiple upstream pathway genes producing strictosidine (MsSLS, MsSTR, Ms7DLGT, MsTDC, MsCPR, Ms8HGO, MsLAMT1, and MsIO). These results provided a basis for identifying candidate enzymes involved in the downstream pathway. Subsequent genomic localization of these three MsMDRs within the published kratom genome^23^ uncovered an MsMDR-enriched BGC spanning approximately 320 kb in Tig00011620 scaffold (Fig. 5b). Consistent with the comparative transcriptomics analysis, this putative BGC contained one gene encoding MsMDR290, three genes encoding MsMDR141 or MsMDR141-like isoenzymes, and four genes encoding known MsDCS1, 3, and 4^24^. Additionally, this BGC contained several additional genes, including genes encoding MsMDR320 and MsMDR4, a functional MDR we identified in a previous study^9^ but which was not reported in the public kratom transcriptome^14^, and multiple uncharacterized MsMDRs and dehydrogenases. In particular, an analog of CpDCE from *Cinchona pubescens*^13^ was also included in this cluster. CpDCE was previously reported to catalyze the hydrolysis of the methyl ester in 20*R*-dihydrocorynantheine (**10a**), the product of CpDCS, which, following spontaneous decarboxylation, produces 20*R*-dihydrocorynantheal (**11a**). The CpDCE analog, designated as MsDCE, showed 85% amino acid sequence similarity and high protein structural similarity with CpDCE^13^ (Supplementary Figs. 14 and 15). Expression levels of MsDCE in published kratom transcriptomes are very low, which likely prevented its earlier identification via expression-based mining. Both CpDCE and MsDCE localized to the yeast cytoplasm as expected (Supplementary Fig. 16), enabling further study of their biosynthetic roles.

### MsDCE and clustered MsMDRs led to novel pathway branches and products

We first investigated the function of MsDCE by co-expressing it with MsSGD and CpDCS in yeast and characterizing the products after 72 hours fermentation supplemented with 0.15 mM strictosidine. CpDCE was included as a control. Like CpDCE, MsDCE converted 20*R*-dihydrocorynantheine (**10a**, *m/z* 355.2017) into 20*R*-dihydrocorynantheal (**11a**, *m/z* of 297.1962), albeit more efficiently, as production of **11a** was ~5 times greater in the assay with MsDCE compared to CpDCE (Fig. 6a). Next we tested whether MsDCE would also accept 20*S*-dihydrocorynantheine (**10b)** as a substrate, which can be produced using MsDCS1 ^9,13,24,25^. We found that MsDCE converts **10a** and **10b** nearly quantitatively into 20*R*- and 20*S*-dihydrocorynantheals (**11a** and **11b**, *m/z* of 297.1962), whereas CpDCE when co-expressed with MsDCS1 only consumed 30.8% of **10a** and 3.40% of **10b** (Fig. 6b). These results suggest that MsDCE could plausibly act as a downstream enzyme of MsDCS1. Co-expressing MsDCE with MsMDR290 and MsSGD in yeast produced a de-esterified compound (*m/z* 295.1805, **12**) from **6** (Fig. 6c), confirming MsDCE’s promiscuity in catalyzing the de-esterification of diverse branches. When co-expressing MsDCE with MsMDR320 and MsSGD in yeast, we only observed **12** derived from **6**; the lack of the corresponding product of **7** is likely due to the lower production of **7** by MsMDR320 as shown in Fig. 4a. Compounds **11a, 11b**, as well as trace amounts of **12** were also detected in kratom leaves (Fig. 6d,e).

## Discussion

We here report the discovery of novel kratom enzymes and their MIA products driven by interactomics analysis using yeast-based PPI screening. We demonstrate that PPIs between different MDRs and the upstream enzyme SGD in MIA biosynthesis are often conserved and can be screened at scale to identify yet unrecognized MsMDR enzymes. Six top-ranked interacting MsMDRs were characterized, revealing enzymatic activity that directly targets the reactive aldehyde form of the strictosidine aglycone (**3**). This chemically labile substrate requires concerted reactions coordinated between MsSGD and MsMDR, which, in turn, explains the strong MsSGD-MsMDR PPIs observed during screening. Genomic analysis using these newly discovered MsMDRs as bait has led to the identification of another type of MIA biosynthesis enzyme, MsDCE, which would have been difficult to predict through transcriptomic analysis due to its extremely low transcript levels in kratom. Overall, integrated pathway prediction efforts, combining interactomics, transcriptomics, and genomics analyses, revealed two pathway branches that produce different de-glycosylated MIAs derived from MsMDRs.

PPIs between SGD and its downstream MDRs^6–9^ are increasingly recognized as critical branching points of MIA biosynthesis across diverse plants, yet there is a lack of systematic understanding of how these PPIs orchestrate MIA biosynthesis within a single plant. Direct interactomics profiling in kratom is limited by technical challenges in plant engineering. As we show here, reconstituting and identifying PPIs in yeast offers an alternative approach to fully profile MsSGD-MsMDR PPIs. Our prior study demonstrated that MsSGD and MsMDRs localize to appropriate organelles in yeast and interact properly^9^, laying the foundation for large-scale screening of the MsMDR library in this work. We also showed that the BiFC assay is a reliable and accurate method for capturing all possible PPIs through the complementation of two split fluorescent protein fragments, which is uniquely suitable for large-scale screening without additional reactions such as split luciferase assays. However, two main obstacles have significantly hindered previous efforts to conduct PPI screening at scale. First, unlike many other types of PPIs successfully screened in yeast or *Arabidopsis*,^26–28^ PPIs between plant MIA enzymes are comparatively weak and transient^29^. Second, a major drawback of the BiFC assay for PPI identification is the high background noise arising from self-assembly of the two split fluorescent protein fragments^30^. As a result, the number of fluorescent proteins resulting from true MsSGD-MsMDR PPIs in most yeast cells is not significantly higher than that in negative control cells, and the overall distribution of cells or average fluorescence in flow cytometry analysis cannot distinguish true interacting pairs from negative controls (Supplementary Fig. 17). To address these challenges, our large-scale PPI screening strategy uses two multi-copy plasmids to express MsSGD and MsMDR in yeast. As the total number of multi-copy plasmids in each yeast cell fluctuates^31^, each yeast construct with positive PPIs always contains a small subpopulation with significantly higher fluorescence than the rest of the cells. This “long-tail” subpopulation is presumably due to higher expression of MsSGD and MsMDR simultaneously, which increases the likelihood of interacting events. Such subpopulations cannot be found or are less common in non-interacting controls and can be used for identifying true PPIs. To further avoid errors in each flow cytometry measurement, we used the subpopulation-to-total population ratios between the candidate pair and the negative control as the criterion to ensure consistent comparisons.

Although the PPI ranks based on the ratio of the high-fluorescent subpopulation cannot precisely quantify their strengths, four of the six newly discovered MsMDRs were ranked at the top of the candidate list. These relatively strong PPIs align with the biochemical function of the MsMDRs. We conclude that interactomics-driven enzyme screening provides a viable strategy for plant natural product biosynthetic pathway discovery, complementing transcriptomics-driven methods^32–35^. However, we note that, although our approach identified all previously known interactors, our interactomics analysis relies on binary PPI screening and cannot detect ternary PPIs in which one protein interacts exclusively at the interface between two other interacting proteins. Further, biochemical screening in yeast in this study focuses on the MsMDRs immediately downstream of MsSGD. Expanding the yeast-based PPI and biochemical screening platform beyond two proteins may enable identification of additional enzymes in MIA biosynthesis.

All previously described MDRs downstream of SGD act on substrates derived from cyclization of the strictosidine aglycone (**3**)^12,36^, whereas the newly discovered MsMDRs act on the very early strictosidine aglycone **3** before cyclization to produce **6**. The instability and cytotoxicity of **3** require rapid enzymatic turnover and controlled transfer, or even metabolite channeling, between MsSGD and MsMDRs. MsMDR290 has shown high activity in yeast for the conversion of **3** into **4**, and the strong PPIs between MsSGD and MsMDR290 or its analogs is consistent with concerted action in planta.

Recent studies have revealed that MDRs in MIA biosynthesis catalyze three types of reactions, including conjugated aldehyde reduction, 1,2-iminium reduction, and 1,4-iminium reduction^12^. The six newly discovered MsMDRs can catalyze the 1,2-reduction of conjugated aldehydes. Different from the other five MsMDRs, MsMDR320 can also catalyze aldehyde reduction of vallesiachotamine (**8**) to vallesiachotaminol (**7**). Production of **7** from **8** was much lower than that of **6** from **3** in both yeast and in vitro assays, likely due to the lower abundance of **8** in our experimental settings and possible competition at binding sites between **8** and **3**. Identification of **7** in kratom stem samples via metabolomics analysis validated it as a natural MIA.

MsMDR141, 120, 262, and 296 did not yield any other novel MIAs besides **6** in yeast when supplemented with strictosidine and co-expressed with MsSGD, based on untargeted metabolomics. However, it remains possible that these four functional MsMDRs, as well as other interacting MsMDRs, may exhibit novel catalytic activities when combined with other kratom enzymes or supplemented with different substrates, as proven by the dual functions of CrDPAS^37^. It is also plausible that enzymes interacting with MsSGD, including MsMDRs and other types of enzymes, may assemble into larger complexes to dynamically regulate MIA biosynthesis. The binary MsSGD-MsMDR PPIs identified, along with yeast-based methods, may provide an entry point for exploring these possibilities.

Kratom metabolomic analysis identified **6, 11**, and a trace amount of **12** in kratom leaves, along with **10** and **11** enriched in stems, suggesting similar pathway branches in planta. This metabolite distribution aligns with the expression levels of corresponding enzymes in different tissues, as MsDCS1, MsMDR290, MsMDR320, and MsMDR141 are all highly expressed in the corresponding tissues. Taken together, the results of our interactomics-driven analysis highlight the potential of integrated multi-omics approaches in identifying plant natural products and their biosynthetic pathways, as well as emphasizing the importance of understanding the spatial organization of enzymes in MIA biosynthesis.

## Methods

### Chemicals and reagents

Yeast nitrogen base (YNB) and amino acid mixtures were purchased from Sunrise Science Products. Ammonium sulfate, dithiothreitol (DTT), tris(2-carboxyethyl) phosphine hydrochloride (TCEP), secologanin, and tryptamine were purchased from SigmaAldrich. Ajmalicine and tetrahydroalstonine were purchased from Neta Scientific. Dextrose, yeast extract (YE), peptone, Luria-Bertani (LB) broth, LB agar, Terrific broth (TB), agar, acetonitrile, formic acid, imidazole, β-nicotinamide adenine dinucleotide phosphate reduced tetrasodium salt (NADPH) and protease inhibitor (100x) were purchased from Thermo Fisher Scientific. All other chemicals, including antibiotics, were purchased from VWR International.

### Microbes and culture conditions

*E. coli* Top 10, *E. coli* BL21(DE3), ccdB resistant *E. coli* and yeast CEN.PK2-1D were used in this work. *E. coli* was used for plasmid construction: Top 10 for normal plasmids and the ccdB resistant strain for plasmids with ccdB gene. All *E. coli* strains harboring plasmids were cultivated in LB media or LB plates at 37°C with 50 μg/mL of kanamycin or 100 μg/mL of carbenicillin as appropriate. Yeast CEN.PK2-1D was used for detecting PPI and studying biochemical functions. Yeasts were cultured at 30°C, 400 rpm in synthetic complete (SC) medium or appropriate synthetic drop-out (SD) liquid media or plates (0.17% YNB, 0.5% ammonium sulfate, 2% dextrose, and amino acid drop-out mixture, and additional 2.5% agar for plates).

### Plasmid construction for gene expression in yeast and *E. coli*

331 MsMDR candidate genes were amplified from plant cDNA. MsDCE and CpDCE were codon-optimized and synthesized by Twist Bioscience. DNA sequences of genes were listed in Supplementary Table 5. Oligonucleotide primers (Supplementary Table 6) were synthesized by Life Technologies. For gene expression in yeast, enzyme-encoding genes were inserted into pre-assembled Gateway-compatible plasmids that contain constitutive promoters and terminators (Supplementary Table 7-8). The gene expression cassettes were assembled into multi-copy expression plasmids using Gateway LR Clonase II Enzyme mix (Life Technologies). For expression in *E. coli*, MsMDR290, MsMDR296 and MsMDR320 were cloned into pET28a plasmid downstream the 6xHis affinity tag using NEBuilder® HiFi DNA Assembly kit (New England Biolabs). Plasmids were extracted using plasmid miniprep kits (Zymo Research), and the sequences were confirmed by Whole plasmid Sequencing (PlasmidSaurus).

### Characterization of PPIs by flow cytometry

Plant cDNA was prepared from 3 μg of each leaf RNA sample via RNA to cDNA EcoDry Premix (Takara Bio) according to the manufacturer’s instructions. After reverse transcription, the product was cleaned with DNA clean kit (Zymo Research) to get around 180 ng of cDNA for each sample.

MsMDR candidate genes were amplified by PCR from kratom cDNA using the Q5 DNA polymerase. BiFC backbones with NV split fragment were also amplified by PCR. PCR products were verified by agarose gel electrophoresis and purified with 96-well DNA cleanup kit from Zymo following the manufacturer’s instructions. Chemically competent yeast cells of the bait strain harboring CV-MsSGD plasmids were prepared and transformed using the standard PEG-LiAc method. 200 ng of backbone with NV split fragment and all purified MsMDR candidate genes were transformed to bait strain. Each well in 96-deep-well plates contained one candidate. Cells after transformation were spread on selection plates SD-WL for 2-4 days of growth.

For BiFC tests, yeast colonies were cultured in deep well plate with 0.5 mL SD-WL at 30°C overnight. 10 μL seed cultures were then diluted in 0.5 mL fresh YPD and cultured for 2 days in duplicates and then used for flow cytometry.

Cell cultures were washed with PBS buffer twice. 20 μL samples were diluted in 180 μL PBS in 96-well plates and measured by flow cytometry. Flow cytometry experiments were conducted in Attune analyzer in biotechnology research center (BRC) at Cornell. The voltage and gate will be settled each time using positive and negative controls, to ensure the ratio of high-fluorescence population (fluorescence >~2000) in MsHYS and MsDCS1 were over 5 folds to the Ps4’OMT negative control, which was less than 0.15%, and that in MsSGD was over 15 folds to the control. Then samples started running with autosampler for 96-well plates. Results were analyzed by FlowJo v10.0.

### In vivo assays of MsMDR candidates and MsDCE using engineered yeast

Plasmids harboring MsSGD and MsMDRs were transformed into yeast CEN.PK2-1D. MsDCE was integrated into the genome by electroporation at a voltage of 540 V, capacitance of 25 μF and infinite resistance with a Gene Pulser Xcell Total System electroporator (Bio-Rad). The correctly colonies were verified through colony PCR analysis. Engineered yeast was cultivated in appropriate SD or SC liquid media or plates at 30°C and tested in triplicates. Strains were cultivated in 96-well plate with 500 µL of SD media at 400 rpm for three days. Strictosidine at a concentration of 0.15 mM was added during fermentation. Cultures were centrifuged at 15000 rpm for 15 minutes, and supernatants were used for LC-MS analysis.

### Characterization of subcellular localization and co-localizations of enzymes in yeast

Plasmids expressing eCFP-MsSGD and mCherry-MsMDRs were individually transformed or co-transformed into the yeast strain CEN.PK2-1D. The strains were cultured in the appropriate SD medium for one day prior to microscopic analysis. Microscopic analysis was performed using a Zeiss LSM 710 Confocal Microscope (AxioObserver, with objective Plan-Apochromat 63X/1.40 Oil DIC M27).

### Protein purification and enzyme assays

All candidates were expressed in *E. coli* BL21(DE3) grown in TB medium. Protein was induced by adding 0.2 mM IPTG and the cultures were shaken at 16°C overnight. Cells were pelleted and lysed by sonication in Buffer A (50 mM Tris-HCl pH 8, 300 mM NaCl, 5% v/v glycerol, 10 mM imidazole), with the addition of an EDTA-free protease inhibitors cocktail. Soluble proteins were purified with Co-NTA magnetic beads (Cube Biotech) and eluted with Buffer B (50 mM Tris-HCl pH 8, 300 mM NaCl, 5% v/v glycerol, 500 mM imidazole). Proteins were concentrated in a 10 kDa cutoff Millipore filter (Merck Millipore) and buffer-exchanged into buffer C (50 mM HEPES pH 7.5). The protein concentrations were measured spectroscopically through MW and extinction coefficients.

The in vitro reaction included 50 mM HEPES pH 7.5, 1 mM strictosidine, 100 mM NaCl, 0.1 mM ZnCl_2_, 1 mM NADPH, 1 μM of the respective MsMDR and MsSGD enzyme. The reaction was incubated at 30°C for 22 hours and quenched by adding 2 volumes of methanol. The mixture was centrifuged at 15000 rpm for 15 minutes, and supernatants were used for LC-MS analysis.

### Strictosidine hydrolysis across pH conditions

To assess the hydrolysis of strictosidine, 0.2 mM strictosidine was incubated in buffers spanning pH 4.0-6.0 (citric acid-sodium citrate buffer), pH 6.0-8.0 (HEPES buffer), and pH 7.0-9.0 (Tris-HCl buffer) for 22 hours. NaCl and ZnCl_2_ were added to approximate in vitro assay conditions. All experiments were performed in triplicates and quenched by adding 1 volume of methanol. The mixture was centrifuged at 15000 rpm for 15 minutes, and supernatants were used for LC-MS analysis.

### Kratom tissue preparation for LC-MS

Kratom tissue samples (~100 mg) were lyophilized for 48 hr using a VirTis BenchTop 4K Freeze Dryer. Lyophilized extracts were suspended in a 4:1 mixture of methanol and water (5 mL) and placed on a shaker overnight at room temperature. The conical tubes containing the extracts were centrifuged (2500 x g, 20 °C, 10 min) and the resulting clarified supernatant was transferred to clean 20 ml scintillation vials, which were concentrated to dryness in an SC250EXP Speedvac Concentrator coupled to an RVT5105 Refrigerated Vapor Trap (Thermo Scientific). The resulting residues were each suspended in 1 ml of a 4:1 mixture of methanol and water, followed by vigorous vortexing and brief sonication. The samples were then transferred to clean microfuge tubes and subjected to centrifugation (10,000 rcf, 20 °C, 5 min) in an Eppendorf 5417R centrifuge to remove precipitate. The resulting supernatants were transferred to HPLC vials without inserts and again concentrated to dryness using the SC250EXP Speedvac Concentrator coupled to an RVT5105 Refrigerated Vapor Trap (Thermo Scientific). The resulting residues were each resuspended in 100 µl of methanol and transferred to clean microfuge tubes and subjected to centrifugation (18,000Xg, 4 °C, 10 min) in an Eppendorf 5417R centrifuge to remove precipitate. The supernatants were transferred to HPLC vials containing inserts and analyzed by LC-HRMS^38,39^.

### LC-MS analysis

Metabolites were analyzed by either an Agilent HPLC-Q-TOF (Agilent 1260 Infinity II/Agilent G6545B) equipped with electrospray ionization source or a Thermo Fischer Scientific Vanquish Horizon UHPLC system coupled with a Thermo Q Exactive HF hybrid quadrupole-orbitrap high resolution mass spectrometer equipped with a HESI ion source. In the Agilent HPLC-Q-TOF, water with 0.1% formic acid (A) and acetonitrile with 0.1% formic acid (B) were used as the mobile phase. For in vitro reaction assays, samples were separated with the Agilent ZORBAX RRHD Eclipse Plus C18 column (2.1 × 50 mm, 1.8 μm), using the gradient elution method of 16 minutes (0-1 min, 95% A; 1-11 min, 95%-5% A; 11-13 min, 5% A; 13-14 min, 5%-95% A; and 14-16 min, 95% A) at 40°C and a flow rate of 0.4 mL/min. For in vivo sample assays, samples were separated with Agilent ZORBAX RRHT Eclipse Plus C18 column (4.6 × 150 mm, 1.8 μm), using the gradient elution method of 60 minutes (0-4 min, 95% A; 4-36 min, 95%-60% A; 36-45 min, 60%-5% A; 45-52 min, 5% A; 52-56 min, 5%-95% A; and 56-60 min, 95% A) at 60°C and a flow rate of 0.6 mL/min. The *m/z* value of the [M+H]^+^ adduct was then used to extract the ion chromatogram (with a mass error below 20 ppm) for compound identification with corresponding chemical standards.

For kratom tissue extracts, LC-HRMS analysis was performed on the Thermo Fischer UHPLC-Q-Orbitrap. HPLC method: water-acetonitrile gradient on an Agilent Eclipse XDB-C18 column (150 mm x 2.1 mm 1.9 µm particle size 175 Å pore size) and maintained at 40 °C. Solvent A: 0.1% formic acid in water; solvent B: 0.1% formic acid in acetonitrile. A/B gradient started at 1% B for 3 min, then from 1% to 99% B linearly over 22 min, then rapidly down from 99% to 1% B and, held at 1% B for 3 min. to equilibrate the column.

To separate charlamine (**6**) and vallesiachotaminol (**7**), an extended 60 min gradient method was applied using the Thermo Fischer UHPLC-Q-Orbitrap. A/B gradient started at 1% B for 3 min, then from 1% to 40% B linearly over 7 min then a more shallowly linear gradient of 40% to 70% B over 45 min, then rapidly down from 70% to 1% B and held at 1% B for 3 min to equilibrate the column.For post-fractionation analysis, water-acetonitrile gradient on an Agilent Eclipse XDB-C18 column (150 mm x 2.1 mm 1.9 µm particle size 175 Å pore size) and maintained at 40 °C. Solvent A: 0.1% formic acid in water; solvent B: 0.1% formic acid in acetonitrile. A/B gradient started at 1% B for 1.5 min, then from 1% to 98% 5.5 min, held at 98% B for 2 min, then rapidly down to 1% B and held for 3 min to equilibrate the column.

Parameters for MS:MS1 resolution, 120,000; AGC target, 3E6; scan range of 130-1000 *m/z*. Parameters for MS/MS (dd-MS2): MS1 resolution, 60,000; AGC target, 3E6; MS2 resolution,30,000; AGC target, 1E5. Maximum injection time, 50 msec; isolation window, 1.0 *m/z*; stepped normalized collision energy (NCE); 30,50; dynamic exclusion, off.

### Nuclear magnetic resonance (NMR) methods

All deuterated solvents were purchased from Cambridge Isotopes. NMR spectra were acquired on a Varian INOVA 600 (600 MHz) spectrometers at Cornell University’s NMR facility at 298.15 K. Alkaloid samples were characterized via a phase-cycled phase-sensitive dqfCOSY spectra using the following parameters: 600 ms acquisition time, 512 complex increments in F1, and 8, 16, 32, or 64 scans per increment, depending on the concentration of the sample. In addition, HSQC, HMBC, and NOESY spectra (using a mixing time of 600 ms), were acquired for all newly identified compounds. All NMR data processing was done using MestreLab MNOVA version 16.0.0-39276.

### Fractionation of MsMDR290 reaction mixture

MsMDR290 reaction mixtures (~40 ml each) containing primarily water and MeOH were diluted with MeOH to 100 ml and added to dry silica (10 g). After rotary evaporation of the solvents the silica was dry-loaded into an empty 25 g Redi*Sep R*f loading cartridge. Fractionation was preformed using a Teledyne ISCO CombiFlash system over a Redi*Sep* Gold 24f HP Silicia column using a dichloromethane-isopropanol solvent system, starting with 95% dichloromethane for 5 column volumes followed by a linear increase of isopropanol content up to 20% over 25 column columns, followed by a sharp linear increase in isopropanol content up to 50% over 2 column volumes, which was then maintained at 50% for an additional 3 column volumes. 30 fractions (16 ml each) generated from the Combiflash run were transferred to clean 20 ml scintillation vials and evaporated *in vacuo* using a SC250EXP Speedvac Concentrator coupled to an RVT5105 Refrigerated Vapor Trap (Thermo Scientific). Dried samples were resuspended in 1 ml of MeOH and subjected to vigorous vortexing and sonication. 50 µl aliquots were transferred to HPLC inserts and subjected to a shorter 12 min gradient that started at 1% B for 1.5 min, then from 1% to 98% B linearly over 5.5 min then held at 98% B for 2 min, then rapidly down from 98% to 1% B and held at 1% B for 3 min to equilibrate the column. LCMS was used for identification of which fractions contained compounds of interest. Fractions containing compounds of interest were pooled together for further analysis by NMR spectroscopy^38^.

### Metabolomics analysis

HPLC-MS HPLC data were analyzed using Metaboseek software^40^ after conversion to mzXML file format using MSConvert. Default Metaboseek settings were used for peak picking and feature grouping. Subsequent detailed analysis of peaks of interest was done using Xcalibur QualBrowser v.4.1.31.9 (Thermo Fisher Scientific). Ion chromatograms were generated using a 5 ppm window around the *m/z* of interest. For isotope pattern simulations a Gaussian profile was used with 200,000 resolution^41^.

### Co-expression analysis of candidates

Hierarchical clustering was performed to visualize the expression patterns of 60 positive interactors and upstream MIA-pathway genes across 49 published kratom transcriptomes^9,14,20–22^, which includes diverse kratom tissues, such as young leaves, mature leaves, leaf bracts, wounded leaves, roots with high or low mitragynine levels, and stems. Transcriptome raw sequence reads used are available from the NCBI Sequence Read Archive under BioProject PRJNA1207432, PRJNA923664, PRJNA664198, PRJNA1114704, PRJNA895092, PRJNA591073, PRJNA849364. Expression levels are presented as Z-scores calculated from log_2_-normalized TPM values. The graph-based clustering heatmap was generated using the R package pheatmap (v1.0.12)^42^.

### Identification of BGC containing MsMDRs and MsDCE

The genomic localization of the MsMDRs was annotated in the kratom genom^23^ on the Tig00011620 scaffold using a custom scrip^43,44^. Then the script searches 320 kbp regions for additional gene annotations by identifying the high sequence similarity with the reported genes and the novel genes discovered in this study. These genes were visualized using the R package gggenomes (v1.1.2)^45^.

### Protein modeling

Protein models were generated using AlphaFold3^46^. Structural alignment between protein MsDCE and protein CpDCE was performed using PyMOL.

### Figure generation

Figures were generated through Prism v10 (Graphpad), FlowJo v10.0, BioRender.com, Adobe Illustrator, Agilent Qualitative Analysis, and Microsoft Office 2016 (PowerPoint, Word, and Excel) whatever necessary.

## Data availability

Genes characterized in this study were deposited to NCBI GenBank under the accession numbers given in Supplementary Table 5. All data are available in the main text or the supplementary materials. Source data are provided with this paper. MS1 data for all samples analyzed in this study will be uploaded to the GNPS web site (massive.ucsd.edu) upon acceptance of the paper.

## Acknowledgments

We thank the Flow Cytometry Facility (RRID: SCR_021740) and the Imaging Facility (RRID: SCR_021741) of the Biotechnology Resource Center of Cornell Institute of Biotechnology. We thank E. Kim, E. Parker Miller, and Y. Zhai for their helpful suggestions and discussions.

## Author Contributions

BY, SK, JH, DH, YW, FS, and SL conceived and designed the research. BY, SK, JH, DH, CL, and YW performed the experiments and analyzed the data. BY, SK, JH, FS, and SL wrote the paper. All authors read and approved the final manuscript.

## Figures

**Extended Data Fig. 1:**
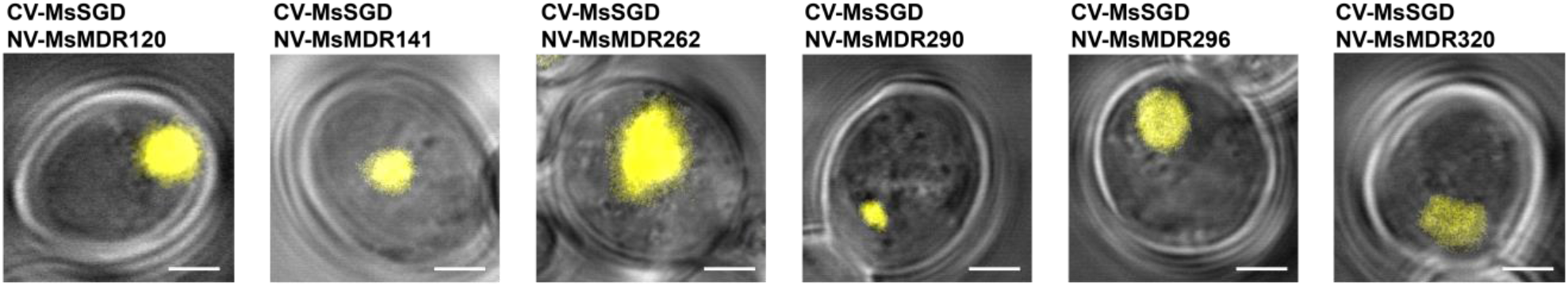
Interactions between MsSGD and six MsMDR candidates in yeast were verified by BiFC. The C-terminal mVenus split fragment (CV) was fused to the N-terminal of MsSGD (CV-MsSGD). And the N-terminal mVenus split fragment (NV) was fused to the N-terminal of functional MsMDRs (NV-MsMDRs). Positive BiFC signals were observed in the yeast nucleus, with fluorescence shown in yellow. Scale bar: 2 μm.

**Extended Data Fig. 2:**
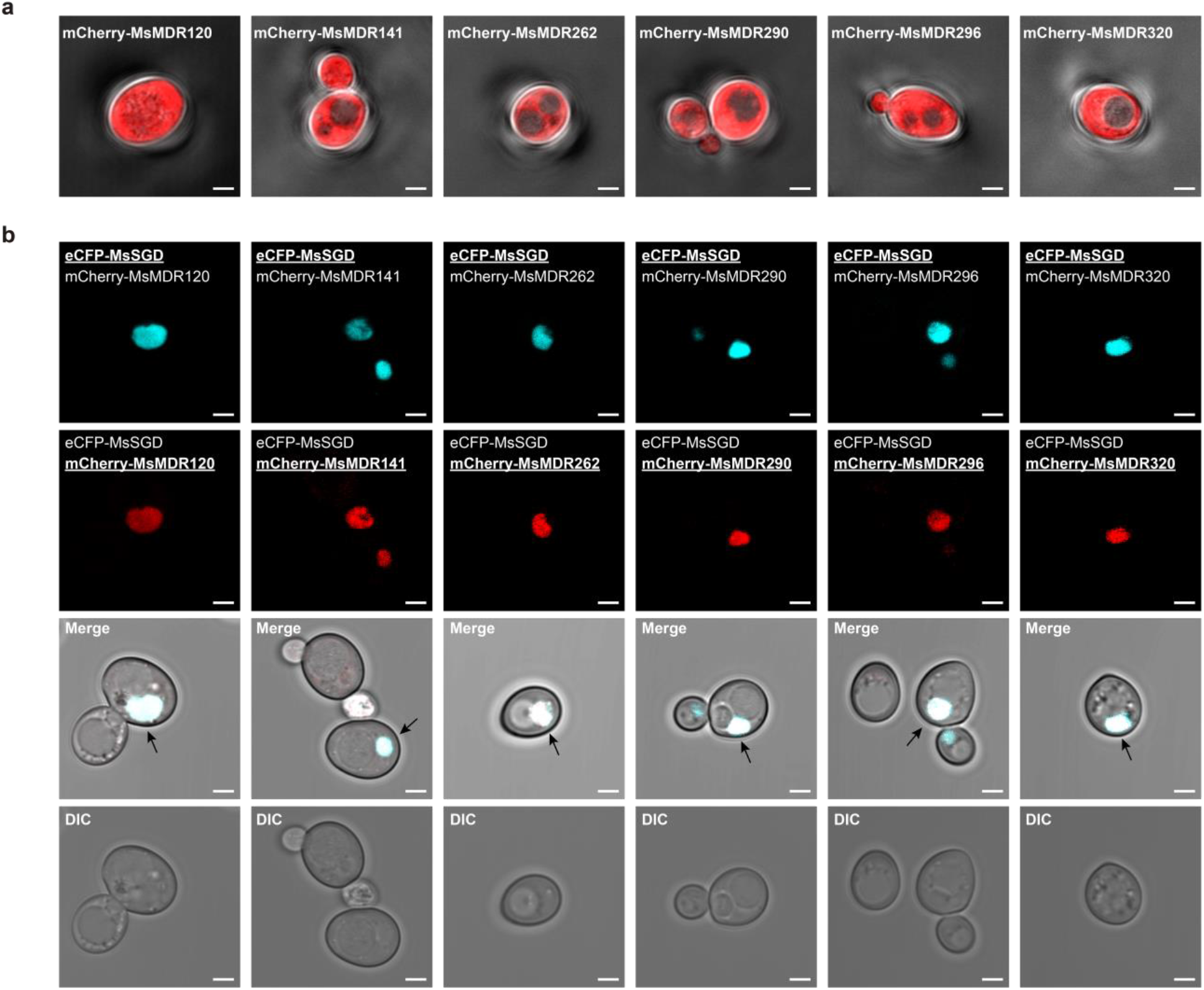
Validated MsSGD-MsMDR interaction in yeast by localization analysis. a. Cytosolic mCherry-tagged MsMDRs (red) when expressed alone. b. MsMDRs (red) localized to the nucleus in the presence of MsSGD (cyan). Merged signals are shown in white and are marked by arrows. Cell morphology was visualized by differential interference contrast (DIC). Scale bar, 2 μm.

